# Potency of nucleoside analogs against Powassan virus replication and identification of 4′-fluorouridine as a therapeutic candidate for tick-borne orthoflavivirus encephalitis

**DOI:** 10.64898/2026.09.24.754293

**Authors:** Devin Shane M. Lewis, Tahirah Moore, Ronak Patel, Robert M. Cox

**Author notes:** contributed equally to this manuscript (Devin Shane M. Lewis, Tahirah Moore).

## Abstract

Powassan virus (POWV) is an emerging tick-borne orthoflavivirus that causes severe encephalitis, long-term neurologic sequelae, and death, yet no approved antiviral therapies exist. Here, we evaluated several previously characterized nucleoside analogs as candidate countermeasures for POWV encephalitis. Favipiravir, molnupiravir, 4′-fluorouridine (4’-FlU), and GS-441524 inhibited POWV replication *in vitro* and demonstrated comparable activity against West Nile virus (WNV). We established C3H/HeNCrl mice as a stringent POWV disease model, showing increased susceptibility, higher day 7 brain titers, and more pronounced neurologic signs resembling severe human disease, compared with C57BL/6 mice. In prophylactic and therapeutic studies, 4′-fluorouridine was the most efficacious compound tested, significantly reducing brain viral burden even when treatment was delayed. Direct comparison with molnupiravir, favipiravir, and GS-441524 in mice confirmed superior survival benefit and antiviral activity for 4′-FlU against both POWV and WNV. These data identify 4′-fluorouridine as a promising therapeutic candidate for neurotropic flavivirus disease.

**Importance:** Powassan virus is an emerging tick-borne orthoflavivirus that causes severe neuroinvasive disease, but there are no approved antivirals for treatment after exposure or onset of neurologic disease. This study addresses that gap by comparing multiple clinically relevant nucleoside analogs against POWV and WNV in cell culture and mouse models of encephalitic orthoflavivirus infection. We identify C3H/HeNCrl mice as a stringent POWV disease model that develops increased brain viral burden and neurologic signs compared with C57BL/6 mice, providing a useful platform for antiviral efficacy studies. Across prophylactic and delayed-treatment experiments, 4′-fluorouridine showed the strongest activity, reducing brain viral titers and improving survival more effectively than favipiravir, molnupiravir, or GS-441524. These findings support 4′-fluorouridine as a lead therapeutic candidate for POWV and WNV and establish a framework for evaluating direct-acting antivirals against neurotropic orthoflavivirus disease.

## Introduction

Powassan virus (POWV) is an emerging tick-borne orthoflavivirus that causes severe neuroinvasive disease in humans and is increasingly recognized as a public health concern in North America. Human infection can range from a mild febrile illness to meningitis, encephalitis, seizures, paralysis, and death. Although reported case numbers remain relatively low compared with mosquito-borne flaviviruses, severe POWV disease is disproportionately neuroinvasive, with an estimated case fatality rate of approximately 10% and long-term neurologic sequelae in a substantial fraction of survivors (1–3). The expanding geographic distribution of Ixodes tick vectors, the rapidity of tick-mediated transmission, and the absence of licensed vaccines or direct-acting antiviral therapies highlight a major gap in preparedness for POWV encephalitis.

POWV comprises two closely related lineages, lineage I and lineage II (deer tick virus), that are serologically indistinguishable and share substantial sequence identity (4–6). Both lineages can cause neuroinvasive disease, and prior work has shown that POWV infection in mice is influenced by viral strain, host genetic background, inoculation route, age, and timing of neuroinvasion (4, 7). These variables have complicated cross-study comparisons and underscore the need for experimentally tractable, immunocompetent mouse models that recapitulate key clinical features of human disease while enabling antiviral efficacy studies. In mice, POWV can invade the central nervous system, produce meningoencephalitis, and establish high viral burdens in the brain; however, susceptibility, kinetics of disease, and clinical penetrance vary by experimental system (8–10). More recent work with C3H-lineage mice has demonstrated robust neurologic disease and uniform lethality following POWV infection, supporting the utility of this host background for modeling severe tick-borne flavivirus encephalitis (11).

Nucleoside and nucleotide analogs represent one of the most clinically validated classes of direct-acting antivirals for RNA virus infections (12–17). Their utility derives from the essential role of the viral RNA-dependent RNA polymerase, the partial conservation of polymerase active-site architecture across related viruses, and the ability of modified nucleosides to interfere with viral RNA synthesis through chain termination, delayed polymerase stalling, nucleotide pool competition, or error catastrophe (13, 18–20). For flaviviruses, this approach is attractive because the NS5 polymerase is indispensable for genome replication and represents a genetically and mechanistically conserved enzymatic target (21–24). Nevertheless, the efficacy of clinically advanced or broadly studied nucleoside analogs has not been systematically evaluated for POWV *in vitro* and *in vivo*.

Here, we evaluated four nucleoside analogs with distinct chemical and mechanistic features, favipiravir, molnupiravir, 4′-fluorouridine (4’-FlU), and GS-441524 (the primary active metabolite of remdesivir), against POWV and West Nile virus (WNV), a related neurotropic orthoflavivirus for which small-animal efficacy benchmarks are better established. We first defined antiviral potency *in vitro* using virus yield reduction assays, assessing potency and ability to reduce progeny virus production. Following this, we then established C3H/HeNCrl mice as a model of POWV encephalitis by comparing disease outcome, clinical presentation, and viral burden with C57BL/6 mice. We next used this model to compare the prophylactic and therapeutic efficacy of each compound. We focused on 4′-FlU, molnupiravir, GS-441524, and favipiravir in mice to further resolve treatment-window effects, survival, and reduction of infectious virus in the brain for POWV and WNV infections. Collectively, these studies identify 4′-FlU as the most active compound in this panel and support further development of ribonucleoside analogs for treatment of tick-borne flavivirus encephalitis.

## Results

### Nucleoside analogs inhibit POWV and WNV replication *in vitro*

We first evaluated the antiviral activity of favipiravir, molnupiravir, 4′-FlU, and GS-441524 against POWV (Lineage II, deer tick virus) and WNV (NY-99 strain) in BHK-21 cells using virus yield reduction assays. Each compound reduced infectious virus production in a dose-dependent manner, although potency differed substantially across the panel (**Figure 1**). All nucleoside analogs inhibited POWV with 50% effective concentrations (EC_50_) values of 10.1 µM, 2.6 µM, 5.1 µM, and 3.3 µM, for favipiravir, EIDD-1931 (the active metabolite of molnupiravir), 4′-FlU, and GS-441524, respectively (**Figure 1A**). In parallel, all nucleoside analogs displayed activity against WNV with EC_50_ values of 30.2 µM, 11.4 µM, 3.1 µM, and 11.2 µM, for favipiravir, molnupiravir, 4′-FlU, and GS-441524, respectively (**Figure 1B**). These data are consistent with the shared dependence of POWV and WNV on the flavivirus NS5 polymerase and support the use of WNV as a comparator virus for benchmarking antiviral efficacy for which prior in vivo POWV data are available. Cytotoxicity was assessed on both BHK-21 cells and primary human astrocytes (**Figure 1C-D**). On BHK-21 cells, cytotoxicity was observed for both EIDD-1931 and GS-441524, with 50% cytotoxic concentrations (CC_50_), respectively. Minimal to no cytotoxicity was observed for both 4’-FlU and favipiravir (CC_50_>100µM). Unlike BHK-21 cells, no cytotoxicity was observed for any nucleoside analog tested on primary human astrocytes. Combined, 4’-FlU exhibited the best performance, with low micromolar potency and no measurable cytotoxic effects.

**Figure 1.**
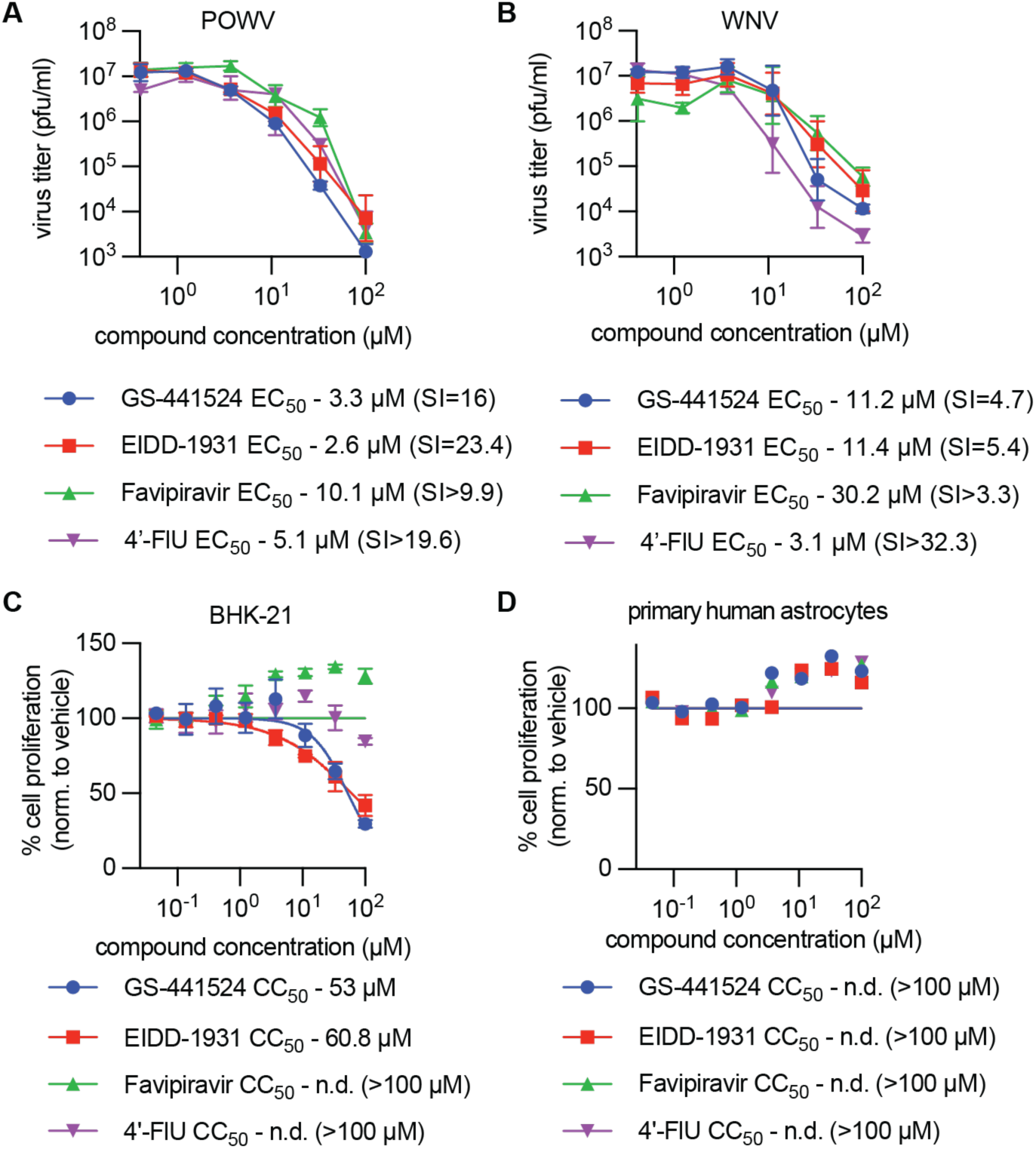
In vitro bioactivity profiles. A-B) Antiviral potency of nucleoside analogs against POWV (DTV) (A) and WNV (NY-99). BHK-21 cells were infected (MOI=0.5) with POWV (A; n=3)) or WNV (B; (n=3)) and incubated with serial dilutions of compound. Supernatant was harvested after 48 hours. Virus titer was determined by plaque assay. **C-D**) Cytotoxicity of nucleoside analogs on BHK-21 (C; (n=3)) and primary human astrocytes (D; (n=3)). Cells were incubated with compound for 72 hours after which effects on cell proliferation and metabolism were determined by PrestoBlue assay. EC_50_ and CC_50_ values were determined by non-linear regression modeling in Prism (version 10.6.1).

### C3H/HeNCrl mice develop severe POWV encephalitis with higher viral burden than C57BL/6 mice

To establish a robust model for antiviral efficacy testing, we compared POWV infection in C3H/HeNCrl using multiple routes of infection and inoculum sizes (**Figure 2A-E**). Regardless of route or inoculum size, all mice succumbed to disease within 13 days. The brains of infected mice displayed an inoculum dependent viral load four days after infection, with animals receiving 10^4^ pfu having an average of 10^6^ pfu per gram tissue, regardless of infection route. Overall animals infected intranasally with 10^2^ and 10^3^ pfu had higher viral loads four days after infection, compared to mice infected subcutaneously (footpad) with equal inoculums. All mice began to show clinical signs by day 6, with prominent clinical signs including bodyweight loss, anxiety (rapid movements, jumping), seizures, and paralysis (**Figure 2C,E**). An inoculum size 1000 pfu was chosen for further study and to establish model criteria for subsequent efficacy studies (**Figure 2F**). For this, we compared intranasal and subcutaneous routes of infection (1000 pfu). Following infection, subgroups of mice (n=3) were sacrificed on days 1, 3, 5, and 7 to assess brain viral loads. Mice infected subcutaneously displayed bodyweight loss earlier than mice infected intranasally (**Figure 2G**). For both routes of infection, neuroinvasion followed a similar trend, with peak titers of ∼10^9^ pfu per gram tissue on day 7. To represent a more natural route of infection, the subcutaneous route of infection with 1000 pfu was chosen for all subsequent POWV efficacy studies.

**Figure 2.**
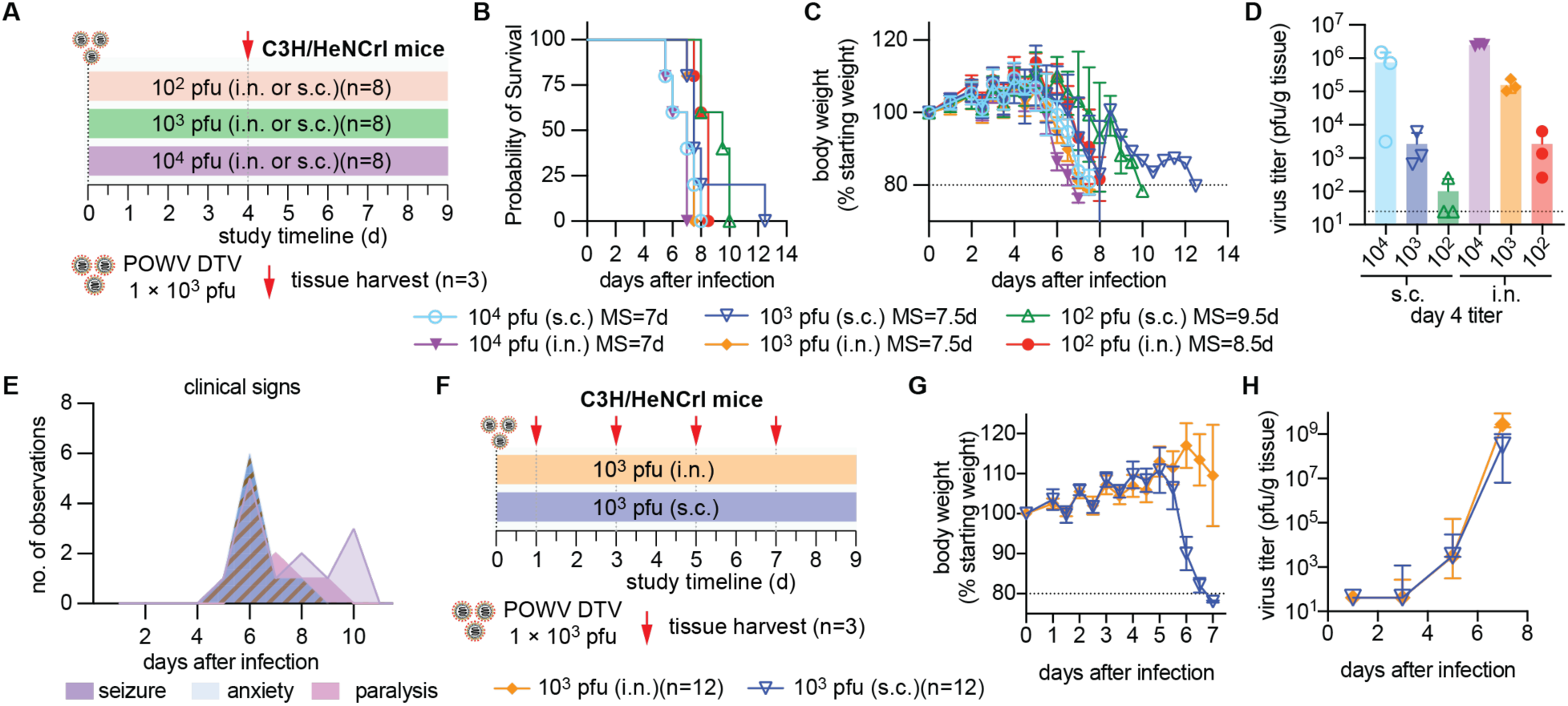
POWV infection in C3H/HeNCrl mice using intranasal and subcutaneous routes of infection. **A**) Study schematic. Groups of C3H/HeNCrl mice (n=8) were infected with different inoculums of POWV using intranasal or subcutaneous routes. Tissue was harvested from subgroups of mice (n=3) on day 4 for virus titration. Animals were monitored daily for clinical signs and bodyweight. **B**) Survival curves. Median survival (MS) in days is shown for each group. **C**) Bodyweight measurements of mice. **D**) Brains of infected mice were harvested 4 days after infection for each route/inoculum (n=3). Virus titer was determined by plaque assay. **E**) Combined summary of major neurological symptoms observed for all groups. All groups displayed similar disease course. F) Study schematic of neuroinvasion following infection with 10^3^ pfu of POWV using intranasal or subcutaneous routes. Mice were infected with 10^3^ pfu of POWV. Subgroups of mice (n=3) were harvested on 1, 3, 5, and 7 days after infection. **G**) Bodyweight measurements **H**) Virus load was determined in the brains mice on days 1, 3, 5, and 7. Lines represent means (C, G) or geometric means (D, H). Errors bars represent standard deviation.

### Prophylactic and Therapeutic efficacy of nucleoside analogs against POWV in C3H/HeNCrl mice

We next compared the prophylactic and therapeutic efficacy of favipiravir, molnupiravir, 4′-FlU, and GS-441524 in the C3H/HeNCrl POWV model (**Figure 3A**). Vehicle mice displayed clinical signs, including bodyweight loss, anxiety, seizures and paralysis, with all animals reaching endpoint by 8 days after infection (**Figure 3B-C**). All treated mice displayed mild to severe body weight loss, regardless of compound, peaking ∼14 days after infection (**Figure 3C**). When treatment was initiated prophylactically, 4′-FlU (5 mg/kg; once daily (*quaque die* (q.d.))) improved survival relative to vehicle-treated animals, whereas favipiravir (150 mg/kg; twice daily (*bis in die*, (b.i.d.))), molnupiravir (150 mg/kg; b.i.d.), and GS-441524 (150 mg/kg; b.i.d.) did not produce a comparable survival benefit (**Figure 3B**). 4’-FlU provided complete protection (100% survival) when administered prophylactically and therapeutically up to 12 hours after infection, and partial survival (60%) when administered 48 h after infection. Although treatment with favipiravir and molnupiravir did not provide complete protection from lethal disease, significant reductions in virus load was observed 7 days after infection (**Figure 3D**). Among the compounds tested, 4′-FlU produced the most consistent virologic benefit and was the only compound that significantly reduced brain virus titers when administered therapeutically, including when treatment was delayed until 48 hours after infection.

**Figure 3.**
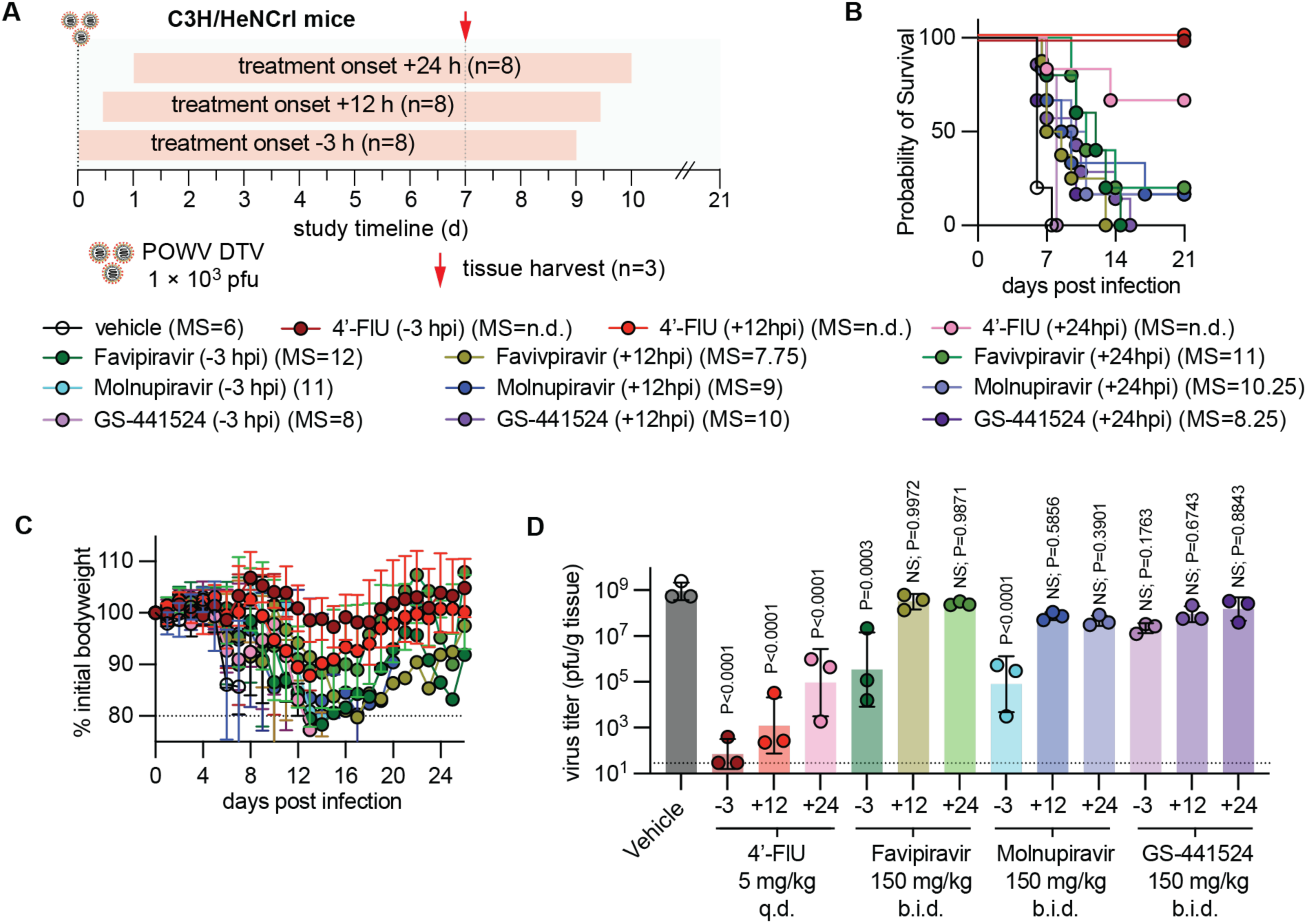
Prophylactic and therapeutic efficacy of nucleoside analogs against POWV in C3H/HeNCrl mice. **A**) Study schematic. C3H/HeN mice (n=8) were infected subcutaneously with 10^3^ pfu of POWV DTV. Mice were treated once daily (q.d.) with 4’-fluorouridine (5 mg/kg) or twice daily (b.i.d.) with either molnupiravir, favipiravir, or GS-441524; starting -3, 12, or 24 hours after infection via oral gavage. Subgroups of mice (n=3) were euthanized 7 days after infection to assess brain virus loads. Bodyweight and clinical signs were monitored daily. **B**) Survival curves. **C**) Bodyweight and clinical signs were monitored daily. Lines intersect means. Symbols represent means. Error bars represent standard deviation. **D**) In subgroups of mice (n=3), brain associated virus loads were determined 7 days after infection. Symbols represent individual repeats. Bars represent geometric means. Error bars represent standard deviation. Statistical analysis was performed using one-way ANOVA with Dunnett’s multiple comparison post test.

### Therapeutic efficacy of nucleoside analogs against POWV in C57BL/6 mice

To confirm efficacy was comparable in other strains, C57BL/6 mice were infected with POWV and treated therapeutically with 4’-FlU (**Figure 4A**). Since numerous other studies have utilized C57BL/6 to study POWV pathogenicity, it was imperative to confirm our findings in this strain (6, 25, 26). Since 4’-FlU was most efficacious in preliminary C3H/HeNCrl studies, it was the only compound selected for assessment in C57BL/6 mice. Treatment was initiated 12, 24, and 48 hours after infection. Compared to C3H/HeNCrl mice, POWV disease was less severe in C57BL/6 mice. Vehicle treated C57BL/6 mice displayed significantly longer time to death and significantly lower viral load on day 7 after infection (**Supplementary Figure 1**)(**Figure 4B-C**). Overall mice displayed similar clinical signs in both C3H/HeNCrl and C57BL/6 mice, with both mice showing weight loss, seizures, paralysis, and death, although C3H appeared to exhibit more anxiety and erratic behavior (jumping in response to stimuli, such as opening cage lids or moving cages). 4′-FlU significantly reduced POWV burden when administered therapeutically, including treatment initiation up to 48 hours post-infection. Treatment starting 12 hours after infection granted complete survival and significant reductions in virus load (**Figure 4B, D**). While treatment starting 24 hours after infection provided for significant reductions in virus titer 7 days after infection, this benefit did not translate to complete survival, with 60% of mice surviving POWV disease. Surprisingly, although later treatment (+48 h) did not lead to significant reductions in virus load, all mice survived infection. These data suggest additional factors, other than reduction in virus load alone, may be involved in survival of POWV infections.

**Figure 4.**
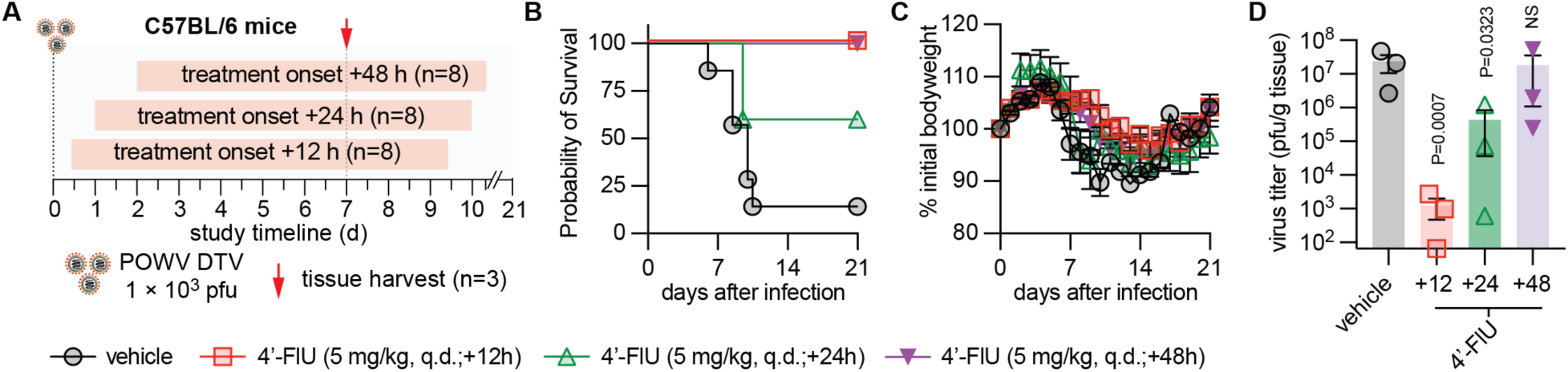
Therapeutic efficacy of 4’-fluorouridine against POWV in C57BL/6 mice. **A**) Study schematic. C57BL/6 mice (n=8) were infected subcutaneously with 10^3^ pfu of POWV DTV. Mice were treated once daily (q.d.) with 4’-fluorouridine (5 mg/kg) starting 12, 24, or 48 hours after infection via oral gavage. Subgroups of mice (n=3) were euthanized 7 days after infection to assess brain virus loads. Bodyweight and clinical signs were monitored daily. **B**) Survival curves. **C**) Bodyweight and clinical signs were monitored daily. Lines intersect means. Symbols represent means. Error bars represent standard deviation. **D**) In subgroups of mice (n=3), brain associated virus loads were determined 7 days after infection. Symbols represent individual repeats. Bars represent geometric means. Error bars represent standard deviation. Statistical analysis was performed using one-way ANOVA with Dunnett’s multiple comparison post test.

### Nucleoside analogs suppress WNV replication in vivo

Given the relatedness of POWV and WNV and the similar *in vitro* potency observed across both viruses, we also evaluated antiviral efficacy in WNV-infected mice. The WNV efficacy studies provide an important comparator because WNV pathogenesis, neuroinvasion, and antiviral response have been more extensively studied than POWV. Prior work comparing C3H and C57BL/6 mice showed that C3H mice are more susceptible to WNV disease than C57BL/6 mice, but that differences in survival are not necessarily explained by major differences in tissue tropism, viral replication kinetics, or CNS viral burden (27).

Initial studies utilized C3H/HeNCrl mice to assess prophylactic efficacy of 4’-FlU, molnupiravir, and favipiravir (**Figure 5**). All mice were infected with 1000 pfu of WNV NY-99 via subcutaneous injection (footpad). All vehicle treated mice reached endpoint by 8 days after infection (**Figure 5B**). Similar to POWV studies, only prophylactic treatment with 4’-FlU provided complete protection from lethal disease. Clinical signs, such as bodyweight loss (**Figure 5C**), were not visible in 4’-FlU treated mice, but all favipiravir treated mice lost weight and succumbed to WNV disease by day 14 following infection. Subgroups of mice were sacrificed 7 days after infection to determine brain viral loads. While all treatments reduced viral loads by 7 days following infection, only 4’-FlU treated mice had significant decreases in viral titers in the brain.

**Figure 5.**
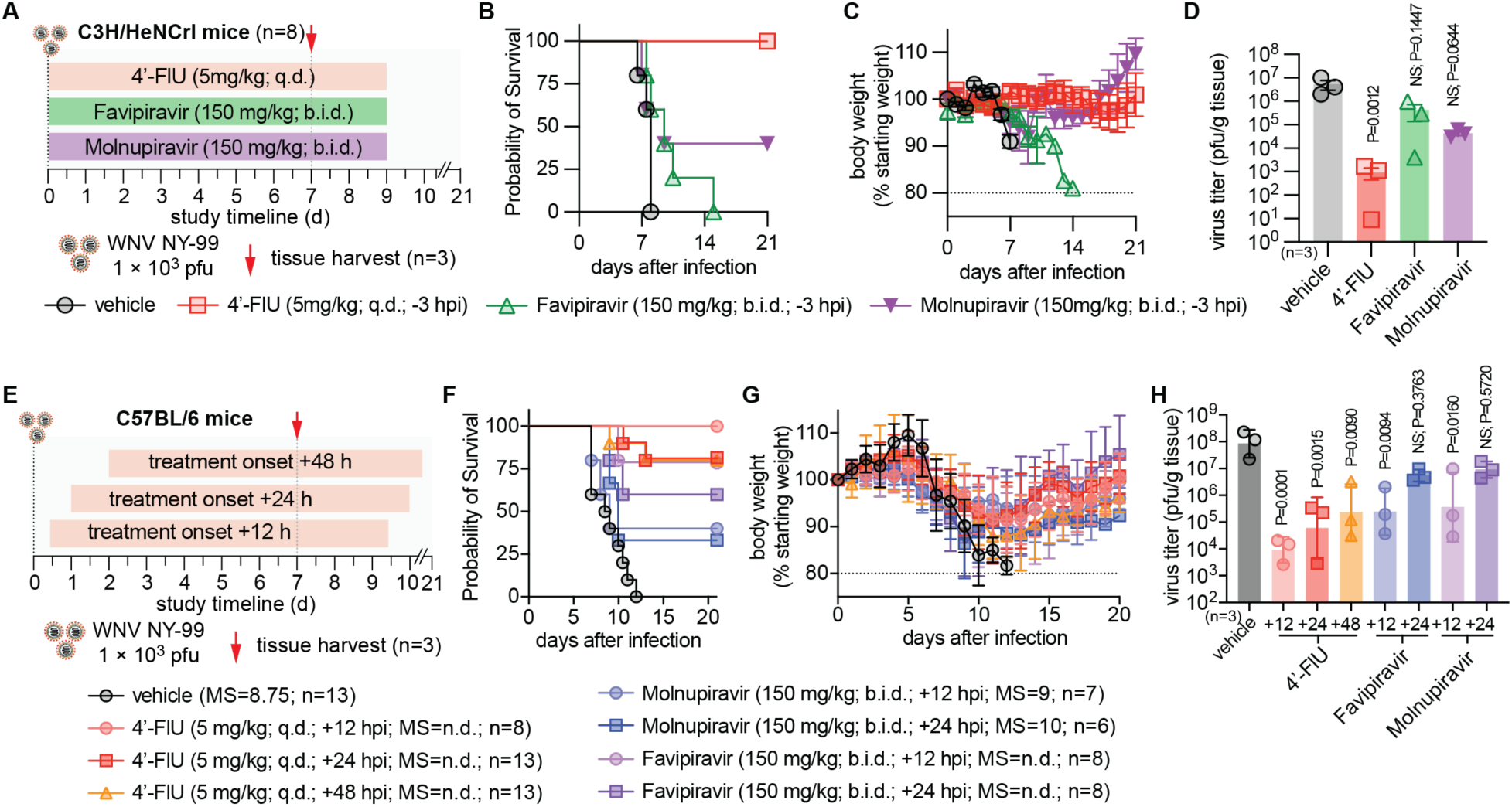
Prophylactic and therapeutic efficacy of nucleoside analogs against WNV NY-99 in C3H/HeNCrl and C57BL/6 mice. **A**) Study schematic. C3H/HeN mice (n=8) were infected subcutaneously with 10^3^ pfu of POWV DTV. Mice were treated once daily (q.d.) with 4’-fluorouridine (5 mg/kg) or twice daily (b.i.d.) with either molnupiravir, or favipiravir; starting 3 hours prior to infection via oral gavage. Subgroups of mice (n=3) were euthanized 7 days after infection to assess brain virus loads. Bodyweight and clinical signs were monitored daily. **B**) Survival curves. **C**) Bodyweight and clinical signs were monitored daily. Lines intersect means. Symbols represent means. Error bars represent standard deviation. **D**) In subgroups of mice (n=3), brain associated virus loads were determined 7 days after infection. Symbols represent individual repeats. Bars represent geometric means. Error bars represent standard deviation. **E**) Study schematic. C57BL/6 mice (n=8) were infected subcutaneously with 10^3^ pfu of POWV DTV. Mice were treated once daily (q.d.) with 4’-fluorouridine (5 mg/kg) or twice daily (b.i.d.) with either molnupiravir or favipiravir; starting 12, 24, and 48 hours after to infection via oral gavage. Subgroups of mice (n=3) were euthanized 7 days after infection to assess brain virus loads. Bodyweight and clinical signs were monitored daily. **F**) Survival curves. **G**) Bodyweight and clinical signs were monitored daily. Lines intersect means. Symbols represent means. Error bars represent standard deviation. **H**) In subgroups of mice (n=3), brain associated virus loads were determined 7 days after infection. Symbols represent individual repeats. Bars represent geometric means. Error bars represent standard deviation. Statistical analysis was performed using one-way ANOVA with Dunnett’s multiple comparison post test. Statistical analysis (D, H) was performed using one-way ANOVA with Dunnett’s multiple comparison post test.

Since previous reports have shown that C3H mice are more susceptible to severe WNV disease, we utilized the less severe C57BL/6 model to further probe the therapeutic potential for 4’-FlU, molnupiravir and favipiravir (27–31). Treatment was initiated 12 or 24 hours after infection, using the same once, or twice daily regimen used in POWV studies (**Figure 5E**). An additional group of 4’-FlU treated mice was included with treatment starting 48 hours after infection. All vehicle treated mice reached endpoint by 12 days after infection (median survival (MS)=8.75 days (**Figure 5F**). All mice experienced transient weight loss, despite outcome of disease (**Figure 5G**). Treatment starting 12 hours after infection improved survival for all treatment groups. Molnupiravir had the least impact on survival, with 40% and 33% survival for +12 (MS=9d) and +24 (MS=10d) hour treatment groups, respectively (**Figure 5F**). Favipiravir treatment had slightly improved survival, with 80% and 60% survival for mice receiving treatment 12 and 24 hours after infection, respectively. 4’-FlU had the greatest impact on survival. All mice receiving 4’-FlU treatment 12 hours after infection survived, and 80% of mice receiving 4’-FlU treatment 24 and 48 hours after infection survived. Treatment with 4’-FlU, molnupiravir and favipiravir 12 hours after infection led to significant reductions in virus titer in the brains of WNV infected mice (**Figure 5H**). However, when treatment was commenced 24 hours after infection, no significant reductions in virus load was observed for mice in with either favipiravir or molnupiravir. Similar to POWV efficacy studies, 4’-FlU treatment led to significant reductions in virus load in all treatment groups (+12, +24, and +48 hours after infection). In our studies, multiple nucleoside analog treatments reduced WNV replication *in vivo*, with 4′-FlU again showing strong antiviral activity. These findings support the broader activity of 4′-FlU against neurotropic flaviviruses and reinforce the utility of POWV and WNV paired efficacy studies for distinguishing compounds with general flavivirus activity from those with virus-specific or model-dependent effects.

## Discussion

This study identifies 4′-FlU as a potent and therapeutically active nucleoside analog for POWV infection and establishes C3H/HeNCrl mice as a rigorous model for evaluating antiviral efficacy against POWV encephalitis. From this study, we have made three major conclusions. First, multiple nucleoside analogs were shown to inhibit POWV replication *in vitro*, confirming nucleoside-based polymerase inhibitors as a valid option for inhibiting POWV replication. Second, we have shown that C3H/HeNCrl mice develop more severe POWV disease than C57BL/6 mice, with clinical manifestations that include seizures, neurologic abnormalities, anxiety-like behavior, weight loss, lethargy, and ataxia, supporting the use of this strain for severe-disease efficacy studies. Third, survival benefit and reduction in CNS viral burden are not equivalent endpoints. Although multiple treatment options could confer significant reductions in virus titer, only 4’-FlU could offer complete protection from lethal POWV disease, even when treatment was initiated up to 12 hours after infection. Direct comparison of 4′-FlU, molnupiravir, and favipiravir in C57BL/6 mice demonstrated that both compounds can reduce POWV burden after infection, but 4′-FlU produced superior survival benefit and retained activity across a wider treatment window.

The *in vitro* potency profile provided an initial rationale for prioritizing 4′-FlU. In BHK-21 cells, 4′-FlU inhibited POWV with an EC_50_ of 5.1 µM. Together with its undefined SI (>19.6), 4’-FlU was identified as the most promising compound tested. Although EIDD-1931 and GS-441524 displayed lower EC_50_ values, their measurable cytotoxicity in the BHK-21 cells used for virus yield reductions may artificially enhance the EC_50_ values recorded. However, *in vitro* potency alone did not predict *in vivo* efficacy. Molnupiravir, favipiravir, GS-441524, and 4′-FlU all showed measurable antiviral activity in cell culture, yet only 4′-FlU consistently reduced POWV titers in the brain and protected from lethal disease when administered therapeutically in mice. This distinction is important because POWV disease is defined by CNS infection and neurologic injury. For neuroinvasive orthoflaviviruses, a clinically meaningful antiviral must do more than inhibit replication in permissive cells; it must achieve sufficient exposure in relevant tissues, suppress viral amplification during the period preceding or accompanying neuroinvasion, ideally reduce the infectious burden in the CNS, and improve disease outcome.

To further characterize *in vivo* efficacy, we compared nucleoside analogs against WNV in C57BL/6 mice using delayed treatment regimens. Three compounds significantly reduced infectious virus titers when administered up to 12 hours after infection, demonstrating that each can suppress POWV replication *in vivo* under conditions where treatment begins after virus exposure. However, the magnitude of protection differed between compounds. 4′-Fluorouridine produced a greater reduction in viral burden and a stronger survival benefit than favipiravir. Complete survival was observed when 4′-fluorouridine treatment began up to 12 hours after infection, and partial survival was retained when treatment was delayed to 24 or 48 hours after infection. In contrast, favipiravir- and molnupiravir-treated mice showed less pronounced reduction in viral load and a greater time-dependent loss of survival benefit.

The C3H/HeNCrl model developed here provides a useful platform for making these distinctions. Prior studies have demonstrated that host genetic background can strongly influence the outcome of orthoflavivirus infection. In WNV, for example, C3H mice exhibit higher morbidity and mortality than C57BL/6 mice despite broadly similar tissue tropism and CNS viral burden, indicating that disease outcome can reflect host-response differences rather than viral replication alone (27). Similarly, POWV pathogenesis is affected by mouse strain, age, virus lineage, route, inoculum, and timing of neuroinvasion (4, 8, 10, 26, 32). Our data show that C3H/HeNCrl mice are more susceptible to severe POWV disease than C57BL/6 mice and develop clinically apparent neurologic signs together with higher viral titers at day 7. This provides a stringent efficacy model in which antiviral activity can be evaluated against a high disease and viral burden background.

The clinical phenotype observed in C3H/HeNCrl mice is notable because human POWV encephalitis is frequently associated with seizures, motor dysfunction, and other neurologic sequelae. Prior mouse studies have shown that POWV infections can produce meningoencephalitis, neuronal injury, gliosis, and viral RNA or antigen in multiple brain regions, but the magnitude and kinetics of disease vary by strain, age, and virus lineage (4, 26, 33, 34). Consistent with these observations, our comparison of C3H/HeNCrl and C57BL/6 mice identified the C3H/HeNCrl background as a more stringent model for antiviral efficacy, particularly for measuring therapeutic effects on CNS viral burden and clinical disease during severe POWV encephalitis.

Among the compounds tested, favipiravir provides the clearest published comparator. Favipiravir has documented activity against WNV, including protection in lethal rodent models when treatment begins shortly after infection. In the original WNV efficacy study, oral T-705/favipiravir administered at 200 mg/kg twice daily beginning 4 hours after subcutaneous WNV challenge protected mice and hamsters from WNV-induced mortality and reduced viral RNA levels in tissues (35). A later mechanistic study showed that favipiravir suppresses WNV in vitro by reducing virus-specific infectivity and increasing mutation frequency, consistent with lethal mutagenesis as at least one mechanism of antiviral activity (28). Those data establish favipiravir as an active anti-WNV compound, but they also emphasize a key limitation: the strongest in vivo efficacy was observed with early treatment initiation. In contrast, our POWV studies show that 4′-fluorouridine retains activity when administered up to 48 hours after infection and provides superior survival benefit compared with favipiravir.

The enhanced efficacy of 4′-FlU is consistent with its broader antiviral profile. 4′-FlU has been shown to be an orally available ribonucleoside analog with activity against multiple RNA viruses, including RSV and SARS-CoV-2, where it acts through inhibition of viral RNA synthesis and polymerase stalling (36). Additional studies have extended its activity to other RNA virus families, supporting the concept that 4′-FlU can function as a broad-spectrum polymerase-directed antiviral scaffold (37, 38). In the present work, its potency against both POWV and WNV, combined with efficacy in two mouse strains and delayed-treatment activity, indicates that 4′-FlU may have utility for neurotropic flavivirus infections where early diagnosis is difficult and therapeutic intervention is likely to occur after systemic infection has begun.

The lack of significant day 7 brain titer reduction by favipiravir, molnupiravir, and GS-441524 in C3H/HeNCrl mice should be interpreted carefully. These results do not necessarily indicate absence of antiviral activity. Favipiravir and molnupiravir prolonged survival when administered prophylactically, suggesting that early suppression of replication, modulation of viral population fitness, or partial reduction in peripheral dissemination may delay disease progression. However, the inability to significantly reduce brain virus titers at the measured time point indicates that these regimens were insufficient to control established CNS-associated POWV burden in this model. The exact nature for the decreased sensitivity *in vivo* of POWV to molnupiravir and favipiravir, whether it relates to mechanism of action, pharmacokinetics, tissue exposure, or polymerase specificity, remains unclear. GS-441524 was active *in vitro* but did not produce a comparable prophylactic survival benefit. The dosing used in the current study was three times higher than previous efficacy studies of HPIV3 in mice, where a dose of only 50 mg/kg b.i.d. was found to be efficacious (39, 40). Our data suggests that either POWV is less susceptible to this compound *in vivo*, that the dosing regimen did not achieve adequate exposure, or that pharmacologic properties limited efficacy in this disease setting. Supporting inadequate exposure, previous studies have shown that administration of GS-441524 in cats did not provide high CNS exposure for the treatment feline infectious peritonitis virus treatments, and additional or higher doses might be required for infections with CNS involvement (41, 42). These findings underscore the importance of integrating pharmacokinetic and tissue exposure data with survival analyses and quantitative virology, particularly in encephalitis models, where delayed mortality, neurologic injury, and persistent viral replication may not closely correlate.

This study has several limitations that can be addressed in future work. First, the present analysis focuses primarily on infectious virus burden and survival, but the relationship between antiviral treatment, CNS inflammation, neuronal injury, and long-term neurologic outcome remains undefined. Because POWV survivors can experience long-term neurologic sequelae, future studies incorporating histopathology, neuroinflammatory markers, behavioral testing, and longer recovery endpoints should be utilized to further probe the effects of treatments on POWV disease and treatment outcome. Second, pharmacokinetic and tissue distribution studies will be needed to determine whether the superior efficacy of 4′-FlU reflects higher systemic exposure, improved tissue penetration, more efficient intracellular triphosphate formation, greater intrinsic potency against POWV polymerase, or a combination of these factors. Finally, susceptibility profiling studies will be required to assess the genetic barrier to escape and to determine whether mutations that confer reduced susceptibility to compound reduce viral fitness or neurovirulence.

The retention of antiviral activity after delayed administration is important given the clinical reality of POWV infection. Tick attachment is frequently unnoticed, early symptoms are nonspecific, and neuroinvasive disease typically emerges only after systemic infection has already been established. Therefore, a compound with activity beyond immediate prophylaxis has substantially greater translational relevance than one that is protective only when administered before or immediately after exposure. These data indicate that 4′-fluorouridine has a broader effective treatment window than the other compounds tested in the C3H/HeNCrl model.

In summary, we establish an immunocompetent C3H/HeNCrl mouse model of severe POWV encephalitis and use it to compare the prophylactic and therapeutic activity of four nucleoside analogs. While multiple compounds inhibited POWV and WNV in vitro, 4′-fluorouridine showed the most favorable in vivo efficacy profile, significantly reducing POWV burden after delayed treatment and providing superior survival benefit relative to favipiravir. These findings separate survival from reduction of CNS viral burden and suggest that the compounds differ not only in intrinsic antiviral potency but also in their ability to achieve the pharmacologic exposure required to suppress POWV replication in vivo. These findings support further development of 4′-fluorouridine as a candidate therapeutic for POWV encephalitis and provide a foundation for broader evaluation of nucleoside analogs against tick-borne and mosquito-borne neurotropic flaviviruses.

## Materials and Methods

### Cells

Baby hamster kidney cells (BHK-21; ATCC) or human primary astrocytes (iXCells Biotechnologies) were maintained at 37°C and 5% CO_2_ in Dulbecco’s modified Eagle’s medium (DMEM) supplemented with 8% fetal bovine serum (FBS) or complete basal astrocyte media (2% FBS with astrocyte growth supplements and pencillin/streptomycin; iXCells Biotechnologies). Immortalized cell lines used in this study are routinely checked for mycoplasma and microbial contamination.

### Molecular virology

POWV DTV stocks were produced from an infectious clone of POWV-DTV (provided by Dr. Margo Brinton, GSU). POWV DTV was rescued under BSL3 conditions. Infectious cDNA clones were linearized and purified by gel extraction or using a Qiagen spin column (Qiagen). Capped RNA transcripts were generated using an mMESSAGE mMACHINE SP6 Transcription Kit. In vitro transcripts were transfected into BHK-21 cells using Lipofectamine MessengerMax transfection reagent, in accordance with the manufacturer’s protocols. Transfected cells were cultured at 37°C for 24-72 hous until CPE was observed. Subsequently, supernatants were then harvested and clarified by centrifugation. Virus stocks were aliquoted and titrated by plaque assay on BHK-21 cells. Prior to virus rescue and for virus stocks, genomes were authenticated using Sanger sequencing.

### Viruses

A WNV NY-99 stock (provided as a gift from Dr. Margo Brinton) was propagated on BHK-21 cells. Virus stocks used in this manuscript were grown and titrated by plaque assay on BHK-21 cells. Virus stocks were authenticated prior to *in vivo* studies.

### Virus yield reduction assay

For virus yield reduction assays, BHK-21 cells were plated 24 h prior (24-well format, 1 × 10^5^ cells per well) and incubated overnight (37°C, 5% CO2). The following day, cells were infected with WNV or POWV (MOI = 0.5) and serial dilutions of nucleoside analogs were added (three-fold dilutions; 100-0.14 µM; plus vehicle-treated). After approximately 48 h, supernatant virus was harvested. Virus titer was determined using plaque assay. Each compound dilution was tested in three independent repeats. Four-parameter variable slope regression modeling was used to determine EC50 concentrations in Prism.

### Cytotoxicity Assays

For cytotoxicity assays, BHK-21 cells or primary human astrocytes (96-well format, 5 × 10^3^ cells per well) cells were plated and incubated overnight (37°C, 5% CO2). The following day, serial dilutions of nucleoside analogs were added (three-fold dilutions; 100-0.045 µM). Cytotoxic concentrations were determined in triplicate after exposure of uninfected cells for 72 h to the different nucleoside analogs. Cytotoxicity was assessed by the addition of PrestoBlue substrate (Invitrogen) to quantify cell metabolic activity, as described (43, 44). Four-parameter variable slope regression modeling was used to determine CC50 concentrations.

### Animal studies

For efficacy studies, C3H/HeNCrl (Charles River) or C57BL/6 (Jackson Labs) mice (6-8 week old; female) were infected with POWV or WNV administered intranasally (1.0 × 10^3^ pfu in 50 μL of PBS) or subcutaneously (via footpad injection (1.0 × 10^3^ pfu in 15µL)). Treatment with 4′-FlU (5 mg/kg; q.d.), molnupiravir (150 mg/kg; b.i.d), favipiravir (150 mg/kg; b.i.d), or GS-441524 (150 mg/kg; b.i.d) commenced -3, 12, 24, or 48 h after infection and continued for 9 days. All analogs were administered by oral gavage in a once daily (q.d.) or twice-daily regimen (b.i.d). Infected mice were monitored twice daily for clinical signs (body weight loss and overall composure) and euthanized upon reaching endpoint or losing >20% of their initial weight. Brain viral titers were determined in groups of mice on day 4 or 7 after infection through plaque assay of brain homogenates on BHK-21 cells. Animal studies were approved under GSU IACUC under protocol A25017.

### Statistics and reproducibility

GraphPad Prism (10.6.1), and Excel (16.92) software packages were used for data analysis. Statistical significance was assessed using ordinary one-way ANOVA with multiple comparisons test in GraphPad Prism, unless otherwise stated. For antiviral potency and cytotoxicity measurements, effective concentrations were determined through four-parameter variable slope regression modeling from dose-response data sets. Biological repeats indicate measurements taken from distinct samples or individual animals.

## Acknowledgements

This work was supported by U19 grant AI171403. Additional thanks to SA for critical reading. The content of this paper is solely the responsibility of the authors and does not necessarily represent the official views of the NIH or NIAID. The funders had no role in study design, data collection and interpretation, or the decision to submit the work for publication.

## Figure Legends

**Supplementary Figure 1.**
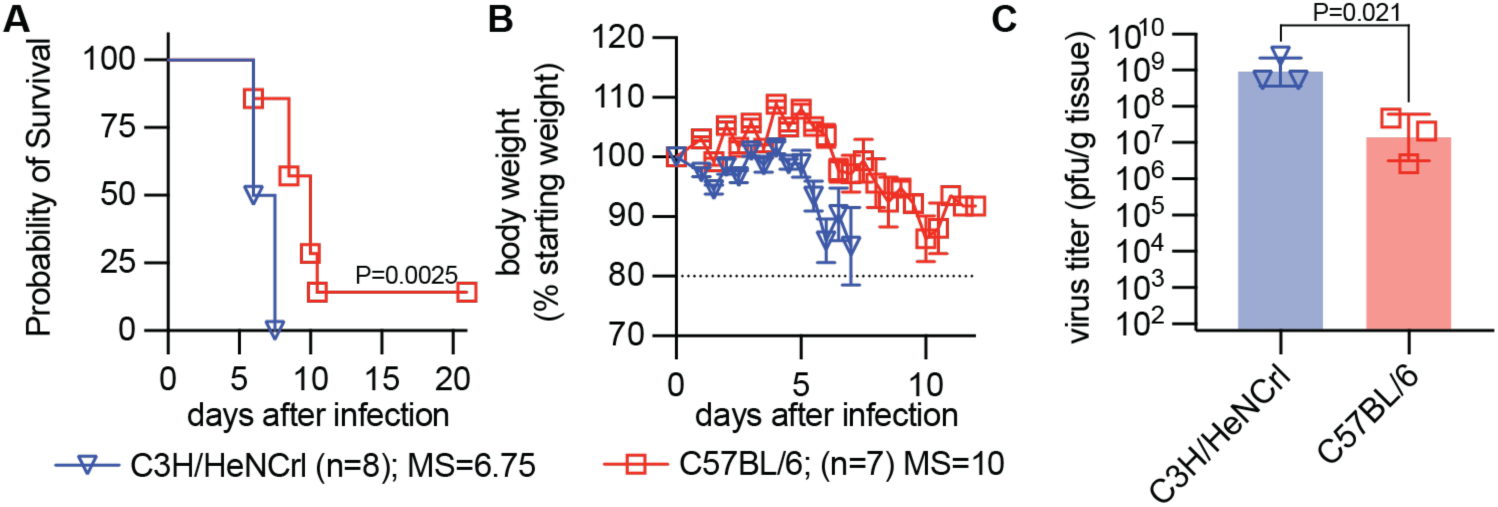
Comparison of POWV infection in C3H/HeNCrl and C57BL/6. **A)** Survival curves of C3H/HeNCrl and C57BL/6 mice as shown in figures 2-4. Survival curves were compared using a Logrank Mantel-Cox test. **B**) Bodyweight measurements as shown in figures 2-4. Lines intersect means. Symbols represent means. Error bars represent standard deviation. **C)** Virus load in the brains of infected mice as shown in figures 3 and 4 for C3H/HeNCrl and C57BL/6 mice, respectively. Symbols represent individual repeats. Bars represent geometric means. Error bars represent standard deviation Statistical analysis was performed using a two-tailed unpaired t test with Welch’s correction.

## References

1. Krow-Lucal ER, Lindsey NP, Fischer M, Hills SL. 2018. Powassan Virus Disease in the United States, 2006-2016. Vector Borne Zoonotic Dis 18:286–290.

2. Hermance ME, Thangamani S. 2017. Powassan Virus: An Emerging Arbovirus of Public Health Concern in North America. Vector Borne Zoonotic Dis 17:453–462.

3. Fatmi SS, Zehra R, Carpenter DO. 2017. Powassan Virus-A New Reemerging Tick-Borne Disease. Front Public Health 5:342.

4. Reynolds ES, Hart CE, Nelson JT, Marzullo BJ, Esterly AT, Paine DN, Crooker J, Massa PT, Thangamani S. 2024. Comparative Pathogenesis of Two Lineages of Powassan Virus Reveals Distinct Clinical Outcome, Neuropathology, and Inflammation. Viruses 16.

5. El Khoury MY, Camargo JF, White JL, Backenson BP, Dupuis AP, 2nd, Escuyer KL, Kramer L, St George K, Chatterjee D, Prusinski M, Wormser GP, Wong SJ. 2013. Potential role of deer tick virus in Powassan encephalitis cases in Lyme disease-endemic areas of New York, U.S.A. Emerg Infect Dis 19:1926–33.

6. Kaur M, Adam M, Mladinich MC. 2025. Pathogenicity and virulence of Powassan virus. Virulence 16:2523887.

7. Courtney SJ, Gallichotte EN, Nilsson E, Trammell CE, Kimball KX, Fagre AC, Vilander A, Overby AK, Ebel GD. 2025. Investigation of Powassan virus lineage II pathogenesis and neurotropism in mice. bioRxiv doi:10.1101/2025.11.07.687151.

8. Jasperse BA, Mattocks MD, Noll KE, Ferris MT, Heise MT, Lazear HM. 2023. Neuroinvasive Flavivirus Pathogenesis Is Restricted by Host Genetic Factors in Collaborative Cross Mice, Independently of Oas1b. J Virol 97:e0071523.

9. Telford SR, Piantadosi AL. 2023. Powassan virus persistence after acute infection. mBio 14:e0071223.

10. Pathak H, Elsharkawy A, Espinola EE, Rothan H, Arora K, Dim C, Stone S, Nabi Z, Kumar M. 2026. Characterization of virus neuroinvasion, blood-brain barrier integrity and neuroinflammation following Powassan virus infection in mice. Front Immunol 17:1813928.

11. Paine DN, Reynolds ES, Hart CE, Crooker J, Thangamani S. 2025. Development of a Deer Tick Virus Infection Model in C3H/HeJ Mice to Mimic Human Clinical Outcomes. Viruses 17.

12. Johnson KA, Dangerfield T. 2021. Mechanisms of inhibition of viral RNA replication by nucleotide analogs. Enzymes 49:39–62.

13. Barik S. 2022. Inhibition of Viral RNA-Dependent RNA Polymerases by Nucleoside Inhibitors: An Illustration of the Unity and Diversity of Mechanisms. Int J Mol Sci 23.

14. Jordan PC, Stevens SK, Deval J. 2018. Nucleosides for the treatment of respiratory RNA virus infections. Antivir Chem Chemother 26:2040206618764483.

15. Stevaert A, Groaz E, Naesens L. 2022. Nucleoside analogs for management of respiratory virus infections: mechanism of action and clinical efficacy. Curr Opin Virol 57:101279.

16. Geraghty RJ, Aliota MT, Bonnac LF. 2021. Broad-Spectrum Antiviral Strategies and Nucleoside Analogues. Viruses 13.

17. Cox RM, Wolf JD, Plemper RK. 2021. Therapeutically administered ribonucleoside analogue MK-4482/EIDD-2801 blocks SARS-CoV-2 transmission in ferrets. Nat Microbiol 6:11–18.

18. Ferrero D, Ferrer-Orta C, Verdaguer N. 2018. Viral RNA-Dependent RNA Polymerases: A Structural Overview. Subcell Biochem 88:39–71.

19. Korneeva VS, Cameron CE. 2007. Structure-function relationships of the viral RNA-dependent RNA polymerase: fidelity, replication speed, and initiation mechanism determined by a residue in the ribose-binding pocket. J Biol Chem 282:16135–45.

20. Campagnola G, McDonald S, Beaucourt S, Vignuzzi M, Peersen OB. 2015. Structure-function relationships underlying the replication fidelity of viral RNA-dependent RNA polymerases. J Virol 89:275–86.

21. Malet H, Masse N, Selisko B, Romette JL, Alvarez K, Guillemot JC, Tolou H, Yap TL, Vasudevan S, Lescar J, Canard B. 2008. The flavivirus polymerase as a target for drug discovery. Antiviral Res 80:23–35.

22. Goh JZH, De Hayr L, Khromykh AA, Slonchak A. 2024. The Flavivirus Non-Structural Protein 5 (NS5): Structure, Functions, and Targeting for Development of Vaccines and Therapeutics. Vaccines (Basel) 12.

23. Lim SP, Noble CG, Shi PY. 2015. The dengue virus NS5 protein as a target for drug discovery. Antiviral Res 119:57–67.

24. Dubankova A, Boura E. 2019. Structure of the yellow fever NS5 protein reveals conserved drug targets shared among flaviviruses. Antiviral Res 169:104536.

25. Davey MG, Riley JS, Andrews A, Tyminski A, Limberis M, Pogoriler JE, Partridge E, Olive A, Hedrick HL, Flake AW, Peranteau WH. 2017. Induction of Immune Tolerance to Foreign Protein via Adeno-Associated Viral Vector Gene Transfer in Mid-Gestation Fetal Sheep. PLoS One 12:e0171132.

26. Courtney SJ, Gallichotte EN, Nilsson E, Trammell CE, Kimball KX, Fagre AC, Vilander AC, Overby AK, Ebel GD. 2026. Pathogenesis and neurotropism of a contemporary Powassan virus lineage II strain in mice. J Virol 100:e0058526.

27. Brown AN, Kent KA, Bennett CJ, Bernard KA. 2007. Tissue tropism and neuroinvasion of West Nile virus do not differ for two mouse strains with different survival rates. Virology 368:422–30.

28. Escribano-Romero E, Jimenez de Oya N, Domingo E, Saiz JC. 2017. Extinction of West Nile Virus by Favipiravir through Lethal Mutagenesis. Antimicrob Agents Chemother 61.

29. Wong J, Moore T, Lewis D, Lieber CM, Vyshenska D, Sobolik EB, Zimmermann S, Painter GR, Plemper RK, Greninger AL, Cox RM. 2026. Impact of mutations affecting 4’-fluorouridine susceptibility on fitness and treatment outcomes for Venezuelan equine encephalitis virus. J Virol 100:e0154125.

30. Ojha D, Hill CS, Zhou S, Evans A, Leung JM, Schneider CA, Amblard F, Woods TA, Schinazi RF, Baric RS, Peterson KE, Swanstrom R. 2024. N4-Hydroxycytidine/molnupiravir inhibits RNA virus-induced encephalitis by producing less fit mutated viruses. PLoS Pathog 20:e1012574.

31. Styer LM, Lim PY, Louie KL, Albright RG, Kramer LD, Bernard KA. 2011. Mosquito saliva causes enhancement of West Nile virus infection in mice. J Virol 85:1517–27.

32. Mladinich MC, Himmler GE, Conde JN, Gorbunova EE, Schutt WR, Sarkar S, Tsirka SA, Kim HK, Mackow ER. 2024. Age-dependent Powassan virus lethality is linked to glial cell activation and divergent neuroinflammatory cytokine responses in a murine model. J Virol 98:e0056024.

33. Scroggs SLP, Offerdahl DK, Stewart PE, Shaia C, Griffin AJ, Bloom ME. 2023. Of Murines and Humans: Modeling Persistent Powassan Disease in C57BL/6 Mice. mBio 14:e0360622.

34. de Souza MRM, Lindner MR, Mladinich-Valenti M, Gorbunova EE, Kirillov V, Byrne AG, Laird AY, Finnerty CE, Ikeda P, Vorkas CK, Kaur R, Gomes YCP, Chatel-Chaix L, Kim HK, Mackow ER. 2026. Single-cell analysis of Powassan virus-infected brains reveals age-dependent neuroinflammatory crosstalk and progressive Alzheimer’s-like APP/Abeta accumulation. mBio 17:e0129426.

35. Morrey JD, Taro BS, Siddharthan V, Wang H, Smee DF, Christensen AJ, Furuta Y. 2008. Efficacy of orally administered T-705 pyrazine analog on lethal West Nile virus infection in rodents. Antiviral Res 80:377–9.

36. Sourimant J, Lieber CM, Aggarwal M, Cox RM, Wolf JD, Yoon JJ, Toots M, Ye C, Sticher Z, Kolykhalov AA, Martinez-Sobrido L, Bluemling GR, Natchus MG, Painter GR, Plemper RK. 2022. 4’-Fluorouridine is an oral antiviral that blocks respiratory syncytial virus and SARS-CoV-2 replication. Science 375:161–167.

37. Lieber CM, Kang HJ, Aggarwal M, Lieberman NA, Sobolik EB, Yoon JJ, Natchus MG, Cox RM, Greninger AL, Plemper RK. 2024. Influenza A virus resistance to 4’-fluorouridine coincides with viral attenuation in vitro and in vivo. PLoS Pathog 20:e1011993.

38. Lieber CM, Plemper RK. 2022. 4’-Fluorouridine Is a Broad-Spectrum Orally Available First-Line Antiviral That May Improve Pandemic Preparedness. DNA Cell Biol 41:699–704.

39. Lin Y, Khan M, Weynand B, Laporte M, Coenjaerts F, Babusis D, Bilello JP, Mombaerts P, Jochmans D, Neyts J. 2024. A robust mouse model of HPIV-3 infection and efficacy of GS-441524 against virus-induced lung pathology. Nat Commun 15:7765.

40. Lin Y, Weynand B, Zhang X, Laporte M, Jochmans D, Neyts J. 2025. The Combination of GS-441524 (Remdesivir) and Ribavirin Results in a Potent Antiviral Effect Against Human Parainfluenza Virus 3 Infection in Human Airway Epithelial Cell Cultures and in a Mouse Infection Model. Viruses 17.

41. Dickinson PJ, Bannasch M, Thomasy SM, Murthy VD, Vernau KM, Liepnieks M, Montgomery E, Knickelbein KE, Murphy B, Pedersen NC. 2020. Antiviral treatment using the adenosine nucleoside analogue GS-441524 in cats with clinically diagnosed neurological feline infectious peritonitis. J Vet Intern Med 34:1587–1593.

42. Gokalsing E, Ferrolho J, Gibson MS, Vilhena H, Anastacio S. 2025. Efficacy of GS-441524 for Feline Infectious Peritonitis: A Systematic Review (2018-2024). Pathogens 14.

43. Cox RM, Sourimant J, Toots M, Yoon JJ, Ikegame S, Govindarajan M, Watkinson RE, Thibault P, Makhsous N, Lin MJ, Marengo JR, Sticher Z, Kolykhalov AA, Natchus MG, Greninger AL, Lee B, Plemper RK. 2020. Orally efficacious broad-spectrum allosteric inhibitor of paramyxovirus polymerase. Nat Microbiol 5:1232–1246.

44. Cox RM, Toots M, Yoon JJ, Sourimant J, Ludeke B, Fearns R, Bourque E, Patti J, Lee E, Vernachio J, Plemper RK. 2018. Development of an allosteric inhibitor class blocking RNA elongation by the respiratory syncytial virus polymerase complex. J Biol Chem 293:16761–16777.

